# In search of non-coding driver mutations by deep sequencing of regulatory elements in colorectal cancer

**DOI:** 10.1101/249300

**Authors:** Rebecca C Poulos, Dilmi Perera, Deborah Packham, Anushi Shah, Caroline Janitz, John E Pimanda, Nicholas Hawkins, Robyn L Ward, Luke B Hesson, Jason WH Wong

**Affiliations:** Prince of Wales Clinical School and Lowy Cancer Research Centre, UNSW Sydney, NSW, Australia; Next-Generation Sequencing Facility, Office of the Deputy Vice-Chancellor (R&D), Western Sydney University, Penrith, NSW, Australia; Department of Haematology, Prince of Wales Hospital, Sydney, NSW, Australia; School of Medical Sciences, UNSW Sydney, NSW, Australia; Faculty of Medicine, The University of Queensland, Herston, QLD, Australia; Level 3 Brian Wilson Chancellery, The University of Queensland, Herston, QLD, Australia; School of Biomedical Sciences, Li Ka Shing Faculty of Medicine, The University of Hong Kong, Hong Kong Special Administrative Region

**Keywords:** colorectal cancer, target capture sequencing, gene regulatory mutations, somatic mutation, mutational signature

## Abstract

Large-scale whole cancer-genome sequencing projects have led to the identification of a handful of cis-regulatory driver mutations in cancer genomes. However, recent studies have demonstrated that very large cancer cohorts will be required in order to identify low frequency non-coding drivers. To further this endeavour, in this study, we performed highdepth sequencing across 95 colorectal cancers and matched normal samples using a unique target capture sequencing (TCS) assay focusing on over 35 megabases of gene regulatory elements. We first assessed coverage and variant detection capability from our TCS data, and compared this with a sample that was additionally whole-genome sequenced (WGS). TCS enabled substantially deeper sequencing and thus we detected 51% more somatic single nucleotide variants (*n* = 2,457) and 144% more somatic insertions and deletions (*n* = 39) by TCS than WGS. Variants obtained from TCS data were suitable for somatic mutational signature detection, enabling us to define the signatures associated with germline deleterious variants in *MSH6* and *MUTYH* in samples within our cohort. Finally, we surveyed regulatory mutations to find putative drivers by assessing variant recurrence and function, identifying some regulatory variants that may influence oncogenesis. Our study demonstrates TCS to be a sequencing-efficient alternative to traditional WGS, enabling improved coverage and variant detection when seeking to identify variants at specific loci among larger cohorts. Interestingly, we found no candidate variants that have a clear driver function, suggesting that regulatory drivers may be rare in a colorectal cancer cohort of this size.

**Author Summary:** In recent years, some cancer research focus has turned towards the role of somatic mutations in the 98% of the genome that is non-coding. To investigate such mutations, we performed deep sequencing of regulatory regions and a selection of coding genes across 95 colorectal cancer and matched-normal samples. To determine the ability of our targeted deep sequencing methodology to accurately detect variants, we compared our results with those from a sample that was additionally whole-genome sequenced. We found target capture sequencing to enable greater sequencing depth, allowing the detection of 51% and 144% more somatic single nucleotide and insertion/deletion mutations, respectively. Our study here demonstrates target capture sequencing to be a useful approach for researchers seeking to identify variants at specific loci among larger cohorts. Our results also enabled the generation of mutational signatures, implicating deleterious germline single nucleotide variants in coding exons of *MSH6* and *MUTYH* in samples within our cohort. Finally, we surveyed regulatory elements in search of somatic cancer driver mutations. We identified some regulatory variants that may influence oncogenesis, but found no candidate variants with clear driver function. These findings suggest that regulatory driver mutations may be rare in a colorectal cancer cohort of this size.

## Background

In recent years, hundreds of novel cancer driver genes have been characterised through analyses made possible by the completion of large-scale cancer-genome sequencing projects. Such genes have been classified as cancer drivers because they harbour frequent high-impact somatic coding mutations in cancer genomes, with these mutations conferring a selective advantage to cells in certain tissues-types and resulting in oncogenesis. Identifying cancer driver mutations outside of protein-coding elements however, has proven to be a complex task as it can be difficult to assign function to some non-coding mutations (1). Despite a number of large-scale studies aimed at prioritising either recurrent or functional mutations (2-4), relatively few somatic driver mutations have yet been discovered in the noncoding genome. One reason for this apparent sparsity of non-coding drivers is that datasets are underpowered to detect mutations at low to moderate frequency from the considerable background of somatic passenger mutations in the cancer genome (5-7).

The costs of whole-genome sequencing (WGS) are constantly decreasing, though performing WGS with sufficient sequencing depth across large cancer cohorts remains expensive. Whole-exome sequencing (WXS) is a potentially cost-effective sequencing alternative for large cohorts, allowing researchers to specifically analyse mutations that arise within protein-coding genes. With the exception of a small proportion of WXS data which can extend into non-target regions including promoter elements (8), WXS cannot identify driver mutations which reside in the remaining ~98% of the genome which is non-coding. In order to refine the potential search area in the non-coding genome, researchers may choose to focus specifically on variants within regulatory elements, such as promoters and other DNase I hypersensitive (DHS) sites which are commonly bound by transcription factors. These sites generally have a greater likelihood of harbouring functional mutations than intergenic regions, as variants at these loci may create or destroy transcription factor binding motifs, or otherwise impact upon nucleosome occupancy or chromatin marks. Recently, sequencing data from an assay capturing regulatory elements in addition to protein-coding regions in a large cohort of breast cancers led to the identification of recurrent somatic mutations in the promoter of the known cancer driver *FOXA1* (5). Therefore, target capture sequencing (TCS) focused on regulatory regions could be an alternative to other sequencing methods, allowing greater cohort sizes, along with increased sequencing depths, at costs comparable to WGS of far fewer samples.

In this study, we perform TCS to generate sequencing data across all promoter elements and some additional regulatory and coding regions in 95 colorectal cancers and matched normal samples. We first assess coverage and variant detection capability from our TCS data, and compare this with a sample that was additionally whole-genome sequenced. We then apply our TCS data to detect mutational signatures, leading to the identification of potentially pathogenic germline variants in patients with suspected sporadic CRC. Finally, we survey somatic mutations in regulatory elements in search of non-coding drivers, finding recurrent somatic mutations in the promoter of *MTERFD3*, as well as some additional variants which may be suitable candidates for further investigation.

## Results

### Target capture sequencing coverage and variant detection

We designed a TCS assay encompassing 35,726,928 nucleotides of the genome (**Fig 1a**; **Table S1a**). The assay was designed to focus on regulatory elements, and primarily covered promoter regions (*n* = 26,455 regions) which we determined using FANTOM5 annotations (9). We also incorporated a selection of DHS sites (*n* = 13,891 regions), long non-coding RNAs (lncRNA; *n* = 842 regions) and microRNAs (miRNA; *n* = 25 regions), at sites where we previously observed mutations in other CRC cohorts. Finally, our panel incorporated coding exons (*n* = 646 exons; *n* = 39 genes) of known colorectal cancer-associated genes (**Table S1b**). With this unique TCS assay, we sequenced 95 colorectal cancer and matched normal samples randomly selected from a pre-existing biobank (**Table 1**; **Table S2**). We obtained high sequencing depth in both cancer and matched normal samples across sequenced regions, with average reads per sequenced base of 169.96 ± 25.08 standard deviation (S.D.) in the cancer samples, and 81.91 ± 17.13 S.D. (S.D. across 95 samples) in the matched normal samples (**Fig 1b**).

**Figure 1.**
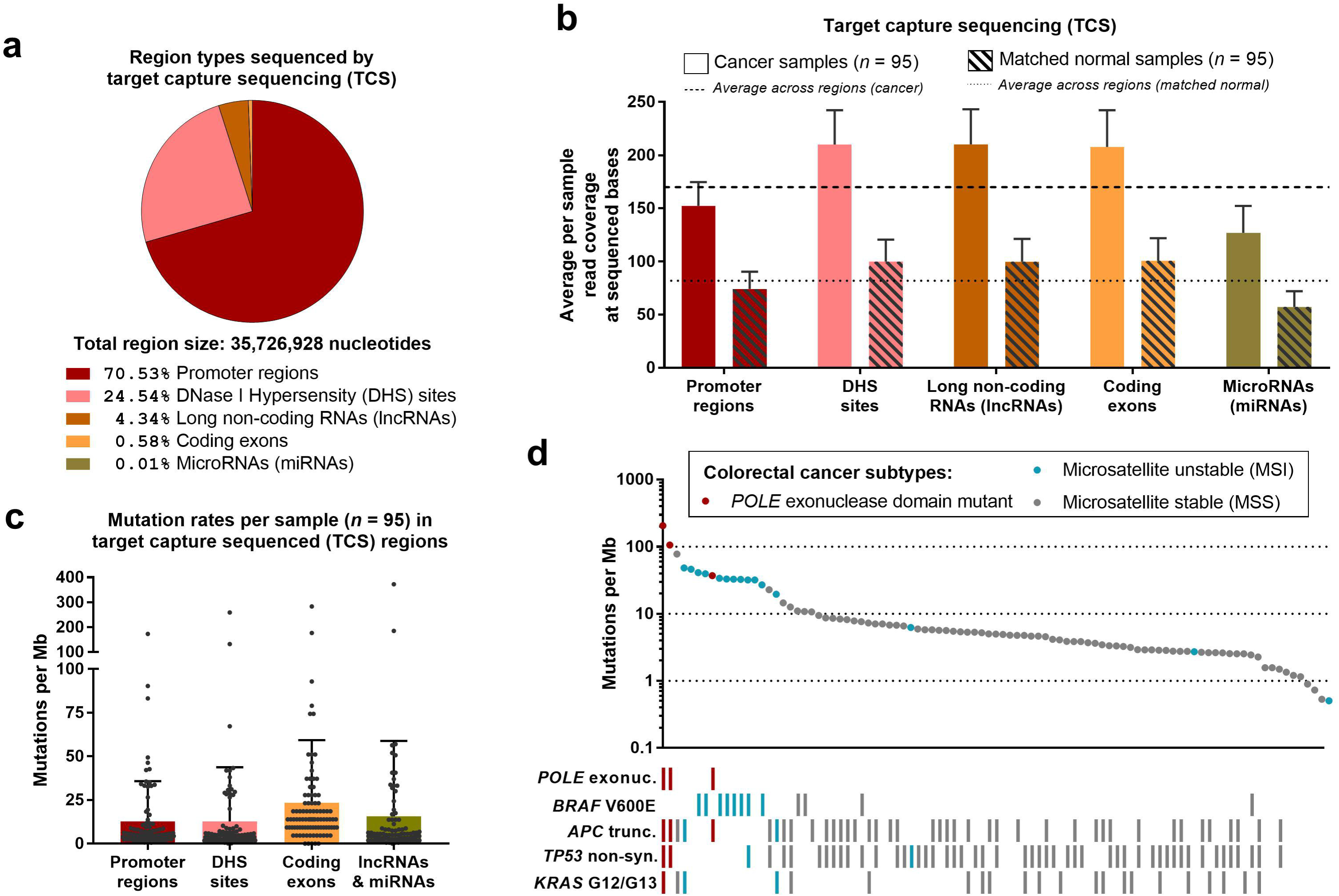
Sequencing coverage by target capture sequencing (TCS), and cohort mutation characteristics. **(a)** Region types sequenced by TCS. Note that 1,107,019 nucleotides of the total region size falls into more than one region type. **(b)** Average per sample reads coverage across sequenced bases in cancer and matched normal TCS samples. Read coverage is plotted for each region type, where box plots show mean and standard deviation across samples in the TCS cohort (*n* = 95). Dotted lines mark average read coverage in tumour and matched normal samples across the cohort. **(c)** Mutations per megabase (mb) in sequenced regions, separated by region type. Dots represent individual samples in the TCS cohort (*n* = 95), and the box plot shows the mean and standard deviation of mutation rates. **(d)** Mutation rate for each individual sample in the TCS cohort (*n* = 95), plotted on a log scale (y-axis). Colours represent individual colorectal cancer subtypes as indicated, and single nucleotide somatic mutations in certain colorectal cancer driver genes are marked by bars. *Exonuc* = exonuclease domain mutation; *trunc* = truncating mutation; *non-syn* = non-synonymous (includes stop gain and stop loss variants).

**Table 1.** Clinicopathological features of the colorectal cancer cohort analysed.

| Characteristic | Cohort (*n* = 95) |
| --- | --- |
| **Age at diagnosis (years; mean ± S.D.)** | 68.8 ± 13.8 |
| **Sex [*n* (%)]** |  |
| Male | 54 (57%) |
| Female | 41 (43%) |
| **Location [*n* (%)]** |  |
| Colon | 53 (56%) |
| Rectum | 42 (44%) |
| **Tumour stage [*n* (%)]** |  |
| Stage I | 31 (33%) |
| Stage II | 32 (34%) |
| Stage III | 32 (34%) |
| **MSI status [*n* (%)]** |  |
| MSI | 15 (16%) |
| MSS | 80 (84%) |
| **CIMP status [*n* (%)]** |  |
| Positive | 12 (13%) |
| Negative | 83 (87%) |
| S.D. = standard deviation; MSI = microsatellite instability; MSS = microsatellite stable; CIMP = CpG Island Methylator Phenotype. |  |

We detected somatic variants using Strelka (10), finding a total of 137,778 single nucleotide somatic mutations within sequenced regions, with a median of 557 somatic mutations per cancer sample. The majority of mutations detected were present at low variant allele frequencies (VAFs; 68% of mutations at ≤ 8.5% VAF). To ensure that we proceeded with analyses of only high-confidence mutation calls, we defined a minimum threshold of ≥ 8.5% VAF to apply to subsequent analyses (**Fig S1a**). Thus, excluding low VAF mutations, the total mutation count across our cohort was 43,915 single nucleotide somatic mutations. Our cancer samples had a median of 178 somatic mutations per sample (**Table S2**), and we validated a selection of somatic single nucleotide mutations, and a deletion mutation via Sanger sequencing (see **Methods**; **Fig S1b-c**).

We observed similar mutation rates at promoters (median 5.02 mutations per megabase [mutations/mb]), DHS sites (4.23 mutations/mb), lncRNA and miRNA (median 5.01 mutations/mb), with coding exons more highly mutated (median 13.93 mutations/mb), consistent with our selection of only known colorectal cancer-associated genes for our TCS assay (**Fig 1c**; raw counts in **Table S2**). By analysing mononucleotide markers as previously described (11), we found 16% (*n* = 15) of our cohort to be microsatellite unstable (MSI). Of the microsatellite stable samples (MSS; *n* = 80), examination of sequencing data revealed that three samples harboured *Polymerase Epsilon (POLE)* exonuclease domain mutations (CRC_1: p.Pro286Arg; CRC_2: p.Met444Lys; CRC_8: p.Ser297Phe), commonly resulting in proofreading deficiency and an ultramutator phenotype (12). Mutation loads across our cohort were generally consistent with previous observations among colorectal cancers (13) (**Fig 1d**), with overall increasing mutation loads in samples which were MSS, MSI and *POLE* exonuclease domain mutated, respectively. Our cohort further reflected known subtype characteristics (14-16), with the MSI samples in our cohort more commonly harbouring *BRAFV600E* mutations (MSI: 8/15; *P* < 0.0001 Fisher’s exact test), and less commonly harbouring *APC* truncating mutations (MSI: 2/15; *P* = 0.0209 Fisher’s exact test), than the *POLE* exonuclease domain wild-type MSS samples (*BRAF V600E*mutation: 4/77; *APC* truncating mutation: 36/77; **Fig 1d**).

### Comparison of target capture and whole-genome sequencing for coverage and variant detection

To assess both the coverage and variant detection capability of our TCS dataset, we selected a sample from our cohort to re-sequence by WGS. We selected the most highly mutated sample in our cohort (CRC_1, with *POLE* exonuclease domain mutation) to ensure that we had large enough numbers of variants for downstream analyses. We performed WGS at the lower depth more commonly associated with this sequencing method, with read coverage averaging 63.61 and 14.29 reads per sequenced base in the cancer and matched normal sample respectively. This coverage was lower than in each of the samples that we sequenced by TCS (**Fig S2a**). In our WGS cancer dataset, the mode coverage was 55-60 reads per sequenced base (11.29%), and 10-15 reads per sequenced base in the matched normal sample (34.79%; **Fig 2b**). In our TCS datasets, the mode coverage was ≥ 100 reads per sequenced base in both the cancer (63.76%) and matched normal (35.34%) samples (**Fig 2b**). When considering coverage in different region types (**Fig S2b-f**), promoters had somewhat low coverage in both TCS and WGS data, likely due to the high GC content in these regions which can lead to poorer sequence coverage and greater rates of misalignment at such loci (5).

**Figure 2.**
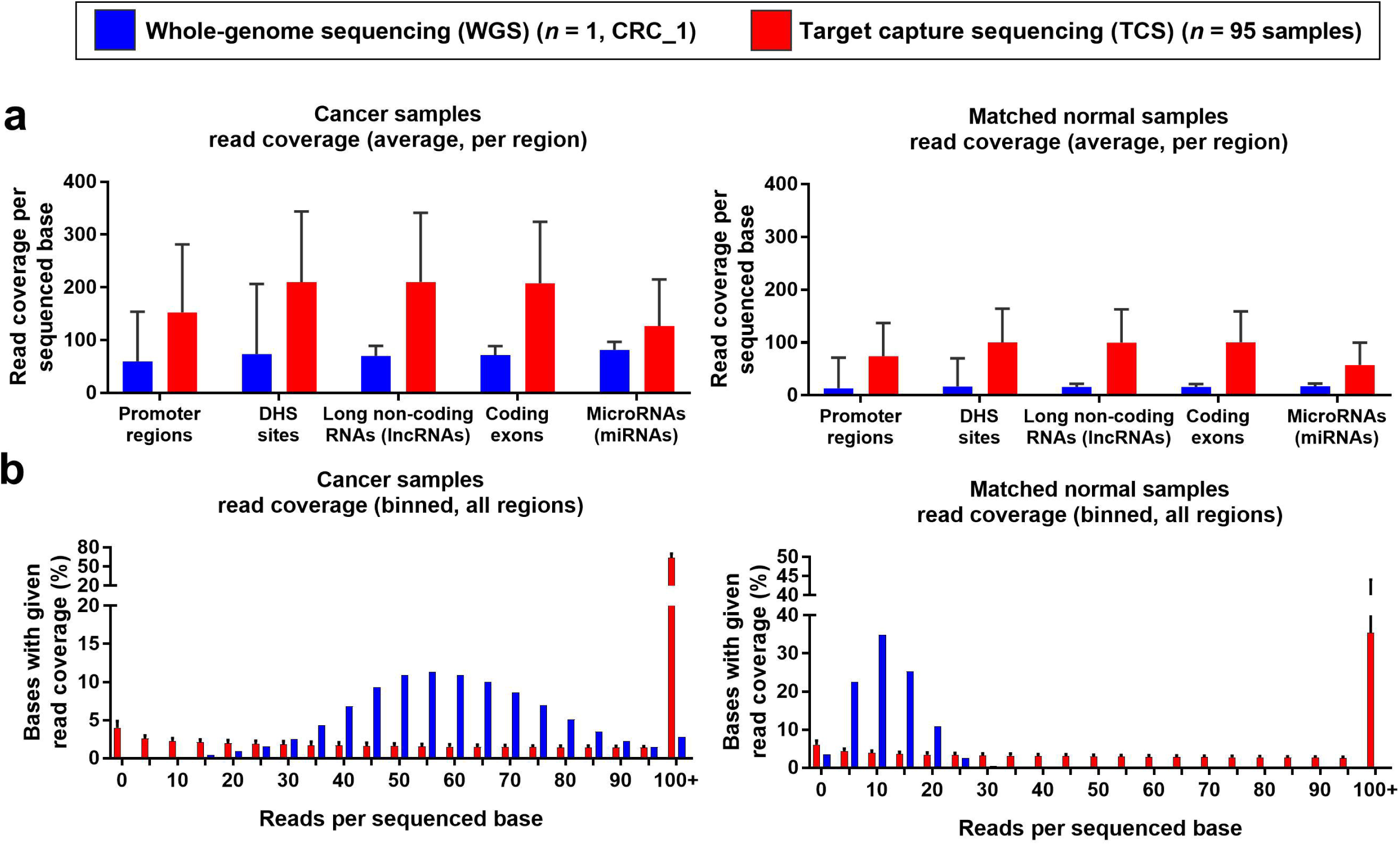
Read coverage statistics for whole-genome sequencing (WGS) and target capture sequencing (TCS) datasets. **(a)** Read coverage per sequenced base in cancer (left) and matched normal (right) samples. Box plot shows mean and standard deviation for all sequenced bases within each region type, where TCS data is pooled across all samples. **(b)** Percentage of bases with given read coverage in cancer (left) and matched normal (right) samples. Data is separated into bins spanning five reads, where the number on the x-axis indicates the lower edge of the bin (inclusive). Box plot shows actual value in WGS data (blue; *n* = 1, CRC_1), and mean and standard deviation across samples in the TCS cohort (red; *n* = 95 samples).

We next compared the somatic mutations that we identified by TCS and WGS, analysing only high-confidence mutations from both datasets (that is, mutations with ≥ 8.5% VAF) within the regions that we incorporated into our TCS assay. We identified 7,311 somatic mutations in CRC_1 via TCS data, but only 4,854 somatic mutations via WGS data (**Fig 3a**). Of these mutations, 4,585 were shared between both TCS and WGS datasets (**Fig 3a**). Interestingly, despite the difference in the absolute numbers of variants detected, the mutational signatures for CRC_1 that were produced using somatic variants from each sequencing method had a Pearson’s correlation coefficient (r) of 0.998 (*P* < 0.0001; **Fig 3b**). Both of these signatures had good correlations with the Catalogue of Somatic Mutations in Cancer (COSMIC) database’s signature 10, which is associated with *POLE* exonuclease domain mutation (TCS: *r* = 0.785 and WGS: *r* = 0.768; *P* < 0.0001; **Fig S3a**). These findings suggest that there was little bias in the trinucleotide composition of the mutations detected by either sequencing method, with the datasets differing primarily in absolute numbers of variants.

**Figure 3.**
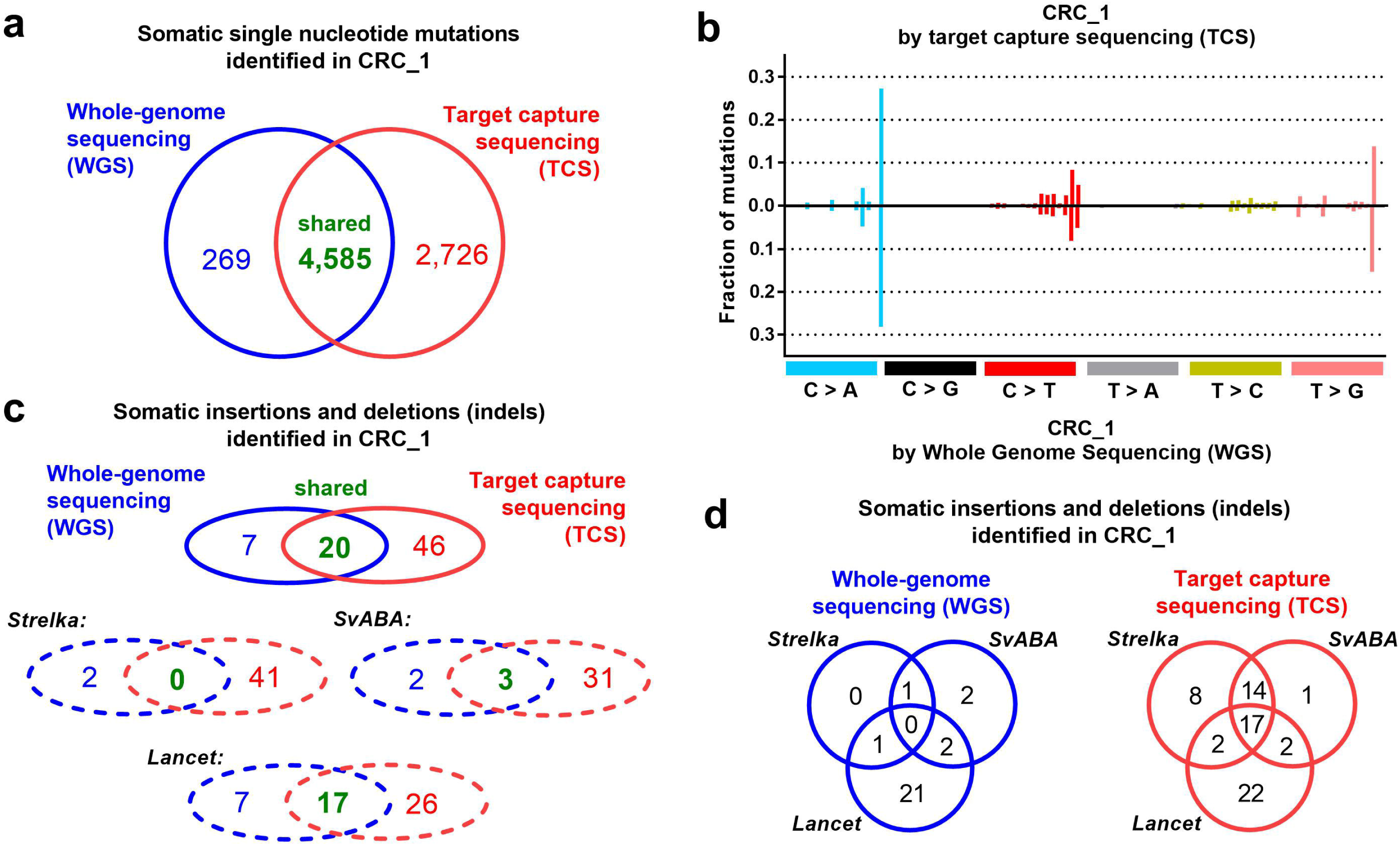
Comparison of variant detection in CRC_1 from whole-genome sequencing (WGS) and target capture sequencing (TCS). **(a)** Venn diagram showing shared and unique single nucleotide somatic mutations identified from WGS and TCS data. **(b)** Mutational signature constructed from single nucleotide somatic mutations identified from TCS (top) and WGS (bottom) data. **(c)** Venn diagram showing numbers of somatic insertions and deletions (indels) identified from WGS and TCS data (solid lines). Venn diagrams indicating numbers of indels identified by different variant detectors are also shown (dotted lines). **(d)** Venn diagrams showing numbers of indels identified by different variant detectors using either WGS or TCS data. All data shown is for colorectal cancer sample CRC_1.

We therefore sought next to investigate why the overall somatic mutation load in CRC_1 differed by TCS or WGS. Hypothesising that low sequencing coverage at some loci might underlie this variation, we examined the sequencing coverage at mutant loci from both TCS and WGS data. Of the 269 somatic mutations that we identified only via WGS data (**Fig 3a**), 47.6% (*n* = 128) had a sequencing depth of ≤ 10 reads in either (or both) of the cancer and matched normal TCS datasets. This is significantly more than in the 4,585 shared somatic mutations identified in both TCS and WGS datasets (0/4,585 [0%]; *P* < 0.0001 Fisher’s exact test). Similarly, of the 2,726 somatic mutations that we identified only via TCS data (**Fig 3a**), 41.6% (*n* = 1,133) had a sequencing depth of ≤ 10 reads in either (or both) of the cancer and matched normal WGS datasets. This too was significantly more than in the 4,585 shared somatic mutations that we identified in both TCS and WGS data (8/4,585 [0.174%]; *P* < 0.0001 Fisher’s exact test). Upon further examination of the sequencing data for the variants with sequencing depth of ≤ 10 reads in WGS data, we found that the low sequencing depth occurred only in sequencing data from the matched normal WGS sample. Of the remaining mutations that had a sequencing depth of > 10 reads in both cancer and matched normal samples, we observed significantly lower coverage at mutant loci in the sequencing dataset from which the mutation was not detected (*P* < 0.0001 unpaired t-test; **Fig S3b-c**). These findings pinpoint sequencing depth as the primary factor underlying the lack of overlap amongst variants detected by the differing sequencing methods. Notably, as matched normal samples are commonly sequenced at lower depths by WGS than the corresponding cancer sample, our study demonstrates this benefit of TCS – which is the increased sequencing depth enabled by focusing only on specific genomic loci.

We next considered the utility of both TCS and WGS for the detection of insertion and deletion (indel) mutations. To do so, we applied three indel callers to both datasets: Strelka (10), SvABA (17) and Lancet (18). Analysing just indels falling into the regions sequenced by our TCS assay, we found that our TCS data enabled the identification of greater numbers of indels (*n* = 66) than did WGS data (*n* = 27, of which 20 indels were shared by both datasets; **Fig 3c**). Lancet detected the highest total number of indels across both samples (*n* = 50), followed by Strelka (*n* = 43) and then SvABA (*n* = 36) (**Fig 3c**). Interestingly, there was very little overlap between the indels identified by all three variant detectors in the WGS data (4/27 [15%] common to two indel callers; 0/27 [0%] common to all three indel callers; Fig 3d). In contrast, in the TCS data, 35/66 indels (53%) were common to at least two indel callers, with 17/66 (26%) identified by all three indel callers (**Fig 3d**). Further, 14/17 (82%) of the indels commonly identified by all three indel callers from the TCS data were among the 20 indels identified by both WGS and TCS. These findings demonstrate TCS to provide greater indel detection sensitivity, and suggest that the variants found from TCS data may be more robust indel calls than those detected by WGS.

### Application of TCS-defined mutational signatures to study cancer pathogenesis

We next investigated cancer pathogenesis via our TCS data through analyses of colorectal cancer subtypes and mutational signatures. We first studied indels detected from our 95 colorectal cancer samples by Strelka (10), SvABA (17) and Lancet (18). We found similar numbers of indels to have been detected by each of the three indel callers (Strelka: *n* = 6,545, SvABA: *n* = 6,603 and Lancet: *n* = 5,649 total indels; **Fig 4a**), with 2,664 indels common to all three variant detectors, and greatest overlap between Strelka and SvABA (total 4,700 indels; Fig 4a). Analysing only high confidence indels detected by at least two of these variant detectors, as expected, we found that MSI samples harboured significantly greater numbers of indels than MSS samples (*P* < 0.0001, unpaired *t*-test; **Fig 4b**). We then defined the mutational signatures for each of the samples in our cohort, using trinucleotide frequencies that have been normalised to match the trinucleotide context of the whole genome. We correlated these signatures with known mutational signatures (19) from the COSMIC database (20, 21). We specifically investigated samples with high correlations with any known signature, in order to assess the utility of TCS for mutational signature analyses.

**Figure 4.**
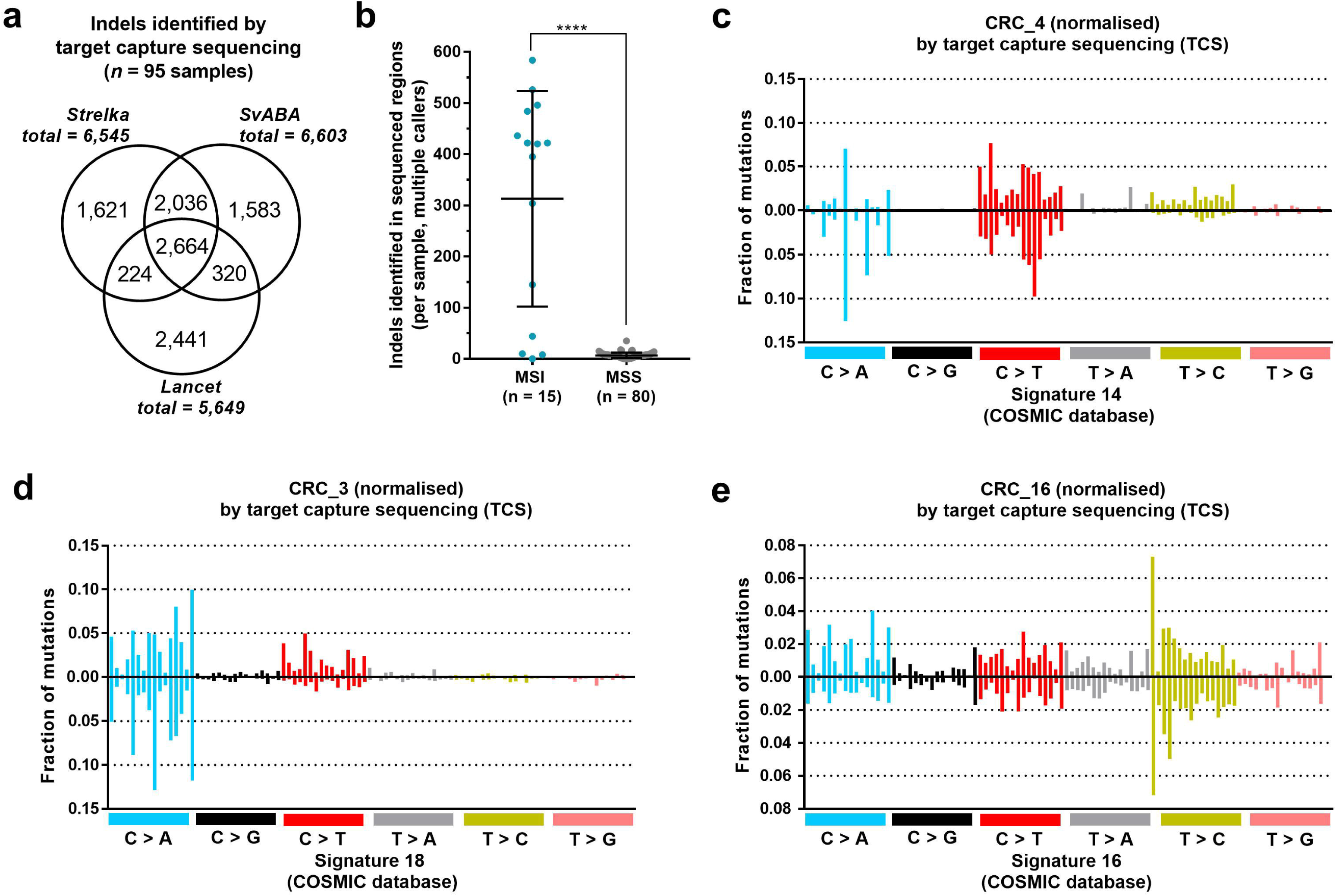
Subtype and mutational signature detection among target capture sequencing (TCS) cohort. **(a)** Total numbers of insertions and deletions (indels) identified by different variant detectors, pooled for the entire TCS cohort (*n* = 95). **(b)** Numbers of indels identified in microsatellite unstable (MSI) and microsatellite stable (MSS) colorectal cancer samples sequenced by TCS. Individual samples are indicated by dots, where counts include indels only identified by at least two different variant detectors. Error bars show mean and standard deviation of indel counts, and **** denotes *P* < 0.0001. (c) Normalised mutational signature from colorectal cancer sample CRC_4 (top), against signature 14 from the COSMIC database (20, 21) (bottom). **(d)** Normalised mutational signature from colorectal cancer sample CRC_3 (top), against signature 18 from the COSMIC database (20, 21) (bottom). (e) Normalised mutational signature from colorectal cancer sample CRC_16 (top), against signature 16 from the COSMIC database (20, 21) (bottom).

We observed strong correlations between the mutational signature of CRC_4 and the COSMIC database’s signatures 14 and 6 (*r* = 0.784 and *r* = 0.767, respectively; and *P* < 0.0001 by Pearson’s correlation; **Fig 4c**). Signature 14 has unknown aetiology but occurs in cancer samples with high mutation loads (19), consistent with CRC_4 being the most highly mutated MSI sample in our cohort (*n* = 1,712 mutations). Signature 6 has been associated with defective mismatch repair and microsatellite instability (19). Given these findings, together with the relatively early age of colorectal cancer diagnosis in this patient (51 years, presenting with synchronous cancers of the rectum and sigmoid), we investigated whether CRC_4 exhibited any germline defects in mismatch repair which might suggest a hereditary cancer predisposition such as Lynch Syndrome. We found CRC_4 to harbour a germline heterozygous C>T SNP at chr2:48,030,588. This SNP is within a coding exon of the mismatch repair gene *MSH6* and results in the introduction of an early stop codon at p.Arg1068* (**Fig S4a**), a variant recorded in the InSiGHT database (22) as Class 5 pathogenic. As a potential somatic second hit that may have contributed to cancer development, CRC_4 also harbours a somatic *MSH6* truncating G>A mutation at chr2:48,026,216 (p.Trp365*). This somatic mutation was present at a VAF of 29%, and loss of MSH6 was evident via immunohistochemistry in both resected tumours. CRC_4 sequencing data exhibited no evidence of *BRAF* V600E mutation, which is additionally consistent with Lynch Syndrome (23) and further supports the results of our mutational signature analysis.

Our mutational signature analyses also highlighted CRC_3 for further investigation, as the mutational signature of this sample was highly correlated with the COSMIC database’s signature 18 (*r* = 0.825 and *P* < 0.0001 by Pearson’s correlation; **Fig 4d**). This sample was the third most highly mutated in our cohort (*n* = 2,767 mutations), which is of particular interest since CRC_3 was neither MSI nor *POLE* exonuclease domain mutated. Signature 18 is characterised by high proportions of C>A variants (19), and has been associated with defects in the base excision repair pathway and *MUTYH* deficiency (24). We found 57% of somatic mutations in CRC_3 to be C>A variants, and so we next examined coding exons of *MUTYH* for deleterious variants. We found no somatic alterations, but instead identified a heterozygous germline C>T SNP at chr1:45,798,117 (**Fig S4b**). This variant has an allele frequency of 1.339x10^-4^ in the Exome Aggregation Consortium (ExAC) database (25), and it causes a non-synonymous amino acid change in *MUTYH* (p.Arg242His) which has been shown *in vitro* to lead to severely defective glycosylase and DNA binding activity (26).

While CRC_3 exhibited no clinicopathological features of MUTYH-Associated Polyposis (MAP), the association with signature 18 suggests that MUTYH alteration by some alternative or additional pathway may have contributed to cancer development in this patient. Our cohort also contained another three samples which had *r* > 0.75 by Pearson’s correlation between their mutational signatures and signature 18 (**Fig S4c**). These samples each had higher mutation loads than the median for MSS samples (median *n* = 162 total mutations), as well as a high proportion of C>A mutations (CRC_19: *n* = 450 total mutations with 50% C>A; CRC_20: *n* = 393 total mutations with 43% C>A; and CRC_26: *n* = 297 total mutations with 53% C>A). However, we found no germline non-synonymous variants in *MUTYH* that were unique to these samples, nor any somatic *MUTYH* mutations. Our findings suggest that these samples may possess larger structural variation affecting *MUTYH* that we are unable to detect via TCS, or that instead some other base excision repair deficiency that would be evident only by examining loci outside of our sequenced regions.

The final signature association that we investigated in detail was between the mutational signature of CRC_16 and the COSMIC database’s signature 16 (r = 0.754 and *P* < 0.0001 by Pearson’s correlation; **Fig 4e**). CRC_16 is a MSS colorectal cancer with a mutation load equivalent to some MSI tumours (*n* = 813 mutations; **Fig 1d**). Recent research suggests that signature 16 in esophageal squamous cell carcinoma may be associated with alcohol intake (27), though signature 16 has primarily been observed in liver cancers and its aetiology remains unconfirmed (19). We found no germline SNPs unique to CRC_16 in any of the exons of the colorectal cancer genes that we sequenced, suggesting that if a germline alteration does explain this signature association, it too may lie outside of our sequenced regions.

In summary, we found that mutational signatures defined only by TCS data that covers a limited portion of the genome can still be sufficient to reveal underlying germline variants involved in cancer pathogenesis.

### Regulatory regions harbouring an excess of functional or recurrent somatic variants

Finally, we sought to identify any regulatory regions that might harbour cancer driver mutations, by examining all somatic single nucleotide and indel variants that we detected from our TCS dataset. To assess the accumulation of functional somatic variants, we applied OncodriveFML (28) to our variants across all sequenced regions listed in **Table S1a**. We first analysed just coding regions of the colorectal cancer driver genes that we sequenced (**Table S1b**), and found many of these genes to be enriched for functional mutations. APC, *KRAS* and *TP53* were the most significantly enriched for functional variants when compared with the expected background mutation load for each gene (**Fig 5a**). In search of regulatory driver mutations, we next excluded coding regions, and used the remaining variants as input for OncodriveFML (28). However, we did not find any regions to be enriched for functional variants in our cohort (**Fig 5b**).

**Figure 5.**
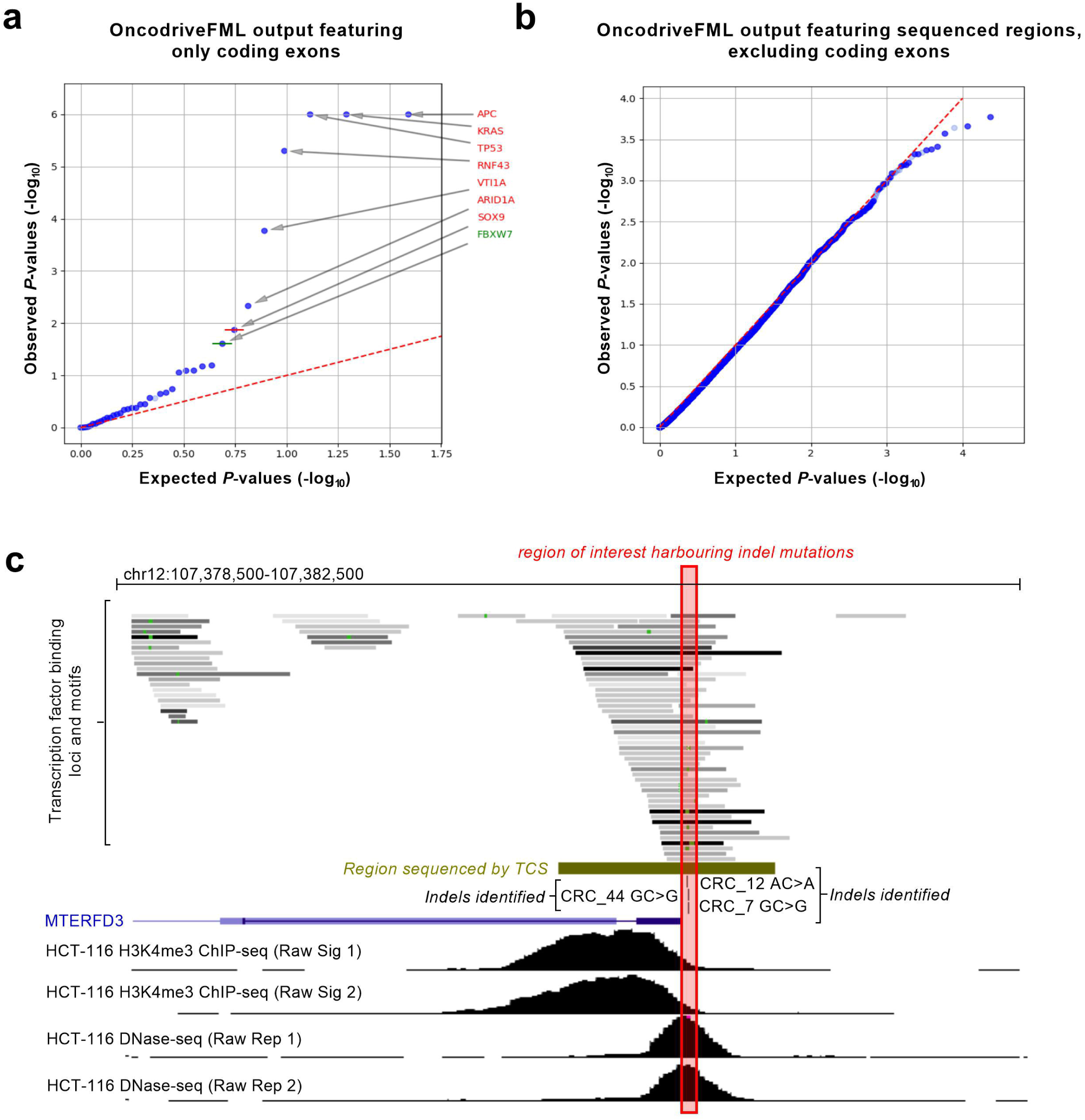
Search for putative driver variants in target capture sequencing (TCS) data. Quantile-quantile plots produced by OncodriveFML (28), showing the expected and observed distribution of functional somatic variant bias P-values **(a)** coding exons of the colorectal cancer-associated genes sequenced and **(b)** all sequenced regions, excluding coding exons from sequenced colorectal cancer-associated genes. Dots represent different sequenced regions, where dots with a lighter colour are regions for which the number of mutated samples did not reach the minimum required to perform the multiple testing correction. Sequenced regions identified as significant are indicated (labels in red: q-value < 0.1 and labels in green: q-value < 0.25). (c) Snapshot from UCSC Genome Browser (51), indicating the location of indels within the putative promoter of *MTERFD3*. Transcription factor binding data is shown via the “Transcription Factor ChIP-seq (161 factors)” track from ENCODE (32). A grey box indicates peak clusters of transcription factor occupancy, where the darkness of each box signifies the maximum signal strength observed in any cell line contributing to that cluster. A green highlight within the box designates the site of the highest scoring canonical motif for the transcription factor indicated, via Factorbook (31) annotations. HCT-116 (human colon cancer cell-line) H3K4me3 chromatin immunoprecipitation sequencing (ChIP-seq) and DNase I hypersensitivity sequencing (DNase-seq) data are also shown.

Assigning function to a non-coding variant can be imprecise due to the variety of ways in which a variant can impact upon gene regulation (1), which can be difficult to capture via a single measure. Hence, in addition to our analyses of functional enrichment in genomic regions via OncodriveFML (28), we also considered base pair recurrence of somatic variants in our cohort. To increase our sample sizes, and to exclude variants which were unique to only our TCS cohort (*n* = 95 samples), we also incorporated single nucleotide variants from WGS colon cancer samples from The Cancer Genome Atlas (TCGA; *n* = 46 samples, **Table S3**) into our analyses. We then selected single nucleotide variants that were present in ≥ 4 samples across cohorts, and at least one TCGA and TCS sample each. Excluding any variants within coding regions of the driver genes that we sequenced, we found 82 recurrent somatic single nucleotide variants (**Table S4**). To prioritise this list for mutations that are more likely to be functional, we annotated these variants using FunSeq2 (29). FunSeq2 annotated 43 of these variants as candidate functional mutations, selected via a high non-coding variant score or an association with any cancer genes (Fu et al., 2014). The 15 mutations with the highest non-coding variant scores are shown in **Table 2**, with the remaining variants listed in **Table S4**. This list of putative functional mutations includes mutations with proximity to cancer related genes such as *JUN, CDKN1B* and *ASF1A* (**Table 2**). The transcription factor binding motif that was most commonly disrupted by the mutations listed in **Table 2** is that for *E2F1* (*n* = 5/15 mutations). The E2F1 protein recognises a binding site consisting of a “CGCGC” DNA sequence (30), in which mutations may more commonly arise as repetitive DNA sequences tend to be more mutagenic.

**Table 2.** Non-coding somatic single nucleotide mutations selected as putative cancer drivers, by base pair recurrence and FunSeq2 annotation.

| Mutation | Recurrence in cohort* |  | % MSI samples | GERP | ENCODE annotation | Motif Analysis |  | Target gene | FunSeq2 Score<sup>^</sup> |
| --- | --- | --- | --- | --- | --- | --- | --- | --- | --- |
|  | TCS | TCGA |  |  |  | Break | Gain |  |  |
| chr7:112,089,887<br>G>T | 3 | 1 | 25% | - | TFP | - | CCNT2_disc2,<br>ZNF740_2,<br>ZNF740_3,<br>ZNF740_4 | IFRD1 (Intron & Promoter) | 4.54 |
| chr1:59,250,792<br>G>A | 3 | 1 | 100% | -0.47 | DHS, TFP,<br>TFM | BCL11A, CCNT2,<br>CTCF, DHS, E2F1,<br>EGR1, ELF1,<br>GABPA, IRF4,<br>MAX, MYC,<br>NFKB1, PAX5,<br>SMARCB1,<br>STAT1, TAF1,<br>TCF12 | - | JUN (Promoter) [**cancer related:** DNA repair, TF regulating known cancer genes, actionable, cancer] | 3.77 |
| chr17:29,648,734<br>A>G | 5 | 1 | 100% | 1.16 | DHS, TFP,<br>TFM,<br>Enhancer | DHS, FOS, MAX,<br>MYC, TBP | HDAC2_disc6 | EVI2A (Promoter & UTR) | 3.75 |
| chr19:41,221,777<br>C>T | 3 | 2 | 100% | -5.51 | DHS, TFP,<br>TFM,<br>Enhancer | DHS, E2F1, ELF1,<br>FOS, FOSL2,<br>GATA2, GTF2F1,<br>JUN, JUND,<br>MAFF, MAFK,<br>NR3C1, RAD21,<br>RFX5, SMARCB1,<br>SMC3, STAT1,<br>STAT3, TCF7L2,<br>USF1, YY1 | - | ADCK4 (Intron & Promoter) | 3.53 |
| chr12:12,957,550<br>T>C | 3 | 1 | 100% | 0.78 | TFP, TFM,<br>Enhancer | EBF1, FOS, JUN,<br>JUND, RFX5,<br>SMARCB1,<br>SMARCC1, TBP | - | APOLD1 (Intron) CDKN1B (Distal) [**cancer related:** DNA repair] DDX47 (Distal) | 3.50 |
| chr4:47,487,706<br>A>T | 3 | 3 | 100% | -5.57 | TFP, TFM | EGR1, MAFK,<br>SPI1 | CPEB1_1 | ATP10D (Intron & Promoter) | 3.25 |
| chr10:104,404,968<br>G>A | 3 | 2 | 60% | -5.86 | TFP, TFM | CTCF, E2F1, TAF1 | - | TRIM8 (Intron & Promoter) | 3.01 |
| chr4:91,049,669<br>C>T | 2 | 2 | 25% | -5.49 | DHS, TFP,<br>TFM,<br>Enhancer | DHS, E2F1, MYC,<br>TCF7L2 | - | CCSER1 (Intron & Promoter) | 2.79 |
| chr11:94,883,648<br>C>T | 2 | 3 | 60% | 0.53 | DHS, TFP,<br>TFM,<br>Enhancer | DHS, E2F1, EP300,<br>FOXA1, GATA1,<br>GATA2, GATA3,<br>MYC | - | - | 2.79 |
| chr11:128,042,476<br>A>G | 3 | 1 | 100% | -7.78 | TFP, TFM,<br>Enhancer | CTCF, FOXA1,<br>RAD21, SMC3,<br>ZNF143 | - | - | 2.79 |
| chr12:4,253,257<br>C>A | 3 | 1 | 50% | -1.65 | DHS, TFP,<br>TFM,<br>Enhancer | CTCF, DHS,<br>RAD21, SETDB1 | - | - | 2.79 |
| chr6:119,215,019<br>A>C | 5 | 1 | 50% | -0.13 | DHS, TFP | - | FOXP1_1 | ASF1A (Promoter) [**cancer**<br>**related:** DNA repair] MCM9<br>(Intron) | 2.78 |
| chr1:154,917,003<br>A>G | 3 | 5 | 88% | - | DHS, TFP,<br>Enhancer | - | - | ADAR (Distal) CKS1B (Distal)<br>EFNA1 (Distal) PBXIP1 (Distal<br>& UTR) PMVK (Distal)<br>PYGO2 (Distal) SHC1 (Distal)<br>ZBTB7B (Distal) | 2.76 |
| chr2:171,787,498<br>A>C | 2 | 2 | 100% | 0.37 | DHS, TFP,<br>TFM | DHS | - | GORASP2 (Intron) | 2.76 |
| chr6:132,272,732<br>A>G | 5 | 1 | 100% | -0.80 | TFP,<br>Enhancer | - | - | CTGF (Promoter) | 2.70 |

\*TCS cohort is the target capture sequencing cohort described in this publication, containing 95 colorectal cancer samples. TCGA cohort is The Cancer Genome Atlas cohort containing 46 whole-genome sequenced colon cancer samples (**Table S3**).
^ This is the “Non-Coding Score” provided by FunSeq2 (29) via a weighted scoring scheme, where higher values indicate variants that may be more likely to be non-coding drivers.
MSI = microsatellite instability. GERP = Genomic Evolutionary Rate Profiling (GERP), a measure of conservation where higher numbers indicate more conserved sites. ENCODE = Encyclopedia of DNA Elements. TFP = transcription factor binding peak; TFM = transcription factor bound motifs in peak region; DHS = DNase I hypersensitive site.

We next investigated recurrent indel mutations, selecting only indels which had been detected by at least two variant detectors for these analyses, as they are less likely to be false positives. We measured indel recurrence within windows spanning 20 base pairs (bp; ±10 bp) so that we could detect regions commonly targeted by indels which can span multiple nucleotides. Analysing only indels in our TCS cohort, we selected genomic windows which harboured ≥ 4 indels, or windows harbouring ≥ 3 indels if at least one of the samples harbouring the recurrent indels was MSS. (Recurrent indels arising in both MSS and MSI samples may be more likely to have arisen because they confer a selective advantage, rather than due to a common mutational process such as microsatellite instability). Excluding indels within coding regions of any of the driver genes that we sequenced, we identified 15 windows ranging in size from 21 bp to 28 bp, which harboured a total of 62 indels (**Table S5**). We sought to prioritise these indels for further investigation by considering their potential impact on transcription factor binding. We ranked indels which lay within transcription factor binding sites, using chromatin immunoprecipitation (ChIP-seq) data and Factorbook annotations (31) from the Encyclopedia of DNA Elements (ENCODE) database (32). The two windows which we found to be the most highly transcription factor-occupied regions were chr15:45,003,769-45,003,795 (indels overlapping a maximum of 71 transcription factor ChIP-seq annotations; *n* = 4 indels) and chr12:107,380,956-107,380,983 (indels overlapping a maximum of 46 transcription factor ChIP-seq annotations; *n* = 3 indels). The former region on chromosome 15 lies within the first exon of *B2M*, and the variants found in our cohort disrupt a repetitive ‘ CTCTCTCTT’ motif within a proteincoding region, and they occurred exclusively in MSI tumours. Indels within exons of *B2M* have already been reported in MSI colorectal cancers, and have been proposed to be involved in colorectal cancer progression (33). Indels in the latter region on chromosome 12 have not previously been described to our knowledge, and we validated all three indels via Sanger sequencing (**Fig S5a**). The region lies within a putative promoter for the mitochondrial transcription termination factor (mTERF) *MTERFD3* (**Fig 5c**). The indels in our cohort overlap Factorbook (31) binding sites for transcription factors *SP1/SP2, E2F4/E2F6* and *MAZ* (**Fig S5b**). Further analysis is limited by the fact that we do not have sample-specific transcriptomic or epigenomic datasets available for each sample in our cohort. However, using data from the colorectal cancer cell line HCT-116, we observed *MTERFD3* expression via RNA sequencing, as well as *SP1* ChIP-seq reads overlapping these indel loci (**Fig S5b**). We also observed *E2F6* and *MAZ* ChIP-seq reads overlapping these indel loci in the HeLa cervical cancer cell line, for which ChIP-seq data in HCT-116 cells were not available (**Fig S5b**). Overexpression of *MTERFD3* and other mTERF family proteins is associated with mitochondrial DNA (mtDNA) copy number depletion (34) and mtDNA copy number variation has been observed in cancer tissues (35). However, experimental functional validation will be required to determine whether these indels might contribute toward oncogenesis through such a capacity.

## Discussion

Over recent years, many recurrent mutations have been identified within *cis-* regulatory regions of cancer genomes, but few drivers have yet been found. This sparsity of non-coding driver mutations may have arisen due to current studies being underpowered to pinpoint drivers present at low to moderate frequencies (5-7). We undertook this study in part to determine whether TCS may enable researchers to increase cohort sizes when seeking to identify driver mutations in defined regions of the genome. We performed WGS at ~60X coverage genome-wide, requiring approximately 900 million 100 bp paired-end reads. Our TCS analyses would have required only 30 million 100 bp paired-end reads per sample (sequencing 35 mb at ~170x), assuming that sequence coverage is only across targeted regions. Therefore, TCS could potentially boost sample sizes by almost 30 fold, whilst also increasing sequencing depth by three fold. By increasing sequencing depth, we identified 51% (*n* = 2,457) and 144% (*n* = 39) more single nucleotide variants and indels, respectively. Therefore, we find TCS to be a sequencing-efficient method to answer specific research questions in large cohorts.

Despite the benefits of TCS that we have demonstrated however, certain limitations upon downstream analyses should be noted from this approach. For example, while we were able to associate CRC_4 and CRC_3 with deleterious germline variants in *MSH6* and *MUTYH* respectively, we were unable to fully investigate the underlying cause of the high mutation load in MSS sample CRC_16, nor the associations that we observed between the additional MSS samples in our cohort and signature 18. The causes of these distinct mutation loads may be a large-scale structural rearrangements, or smaller variants in other regions of the genome, that we were unable to investigate without further sequencing. TCS is likely to be unsuitable for such investigations of a more exploratory nature where researchers may need to extend analyses into regions of the genome not initially included in a TCS assay. Further, some non-coding driver mutations create *de novo* promoter and enhancer regions affecting important cancer-associated genes (36-38). Therefore, another limitation of TCS for non-coding driver detection is that any somatically-acquired regulatory regions that harbour driver mutations could remain undetected, as these regions may not have been selected for inclusion into a TCS assay. This limitation applies to this current study, as the DHS regions sequenced were selected using only a single colorectal cancer cell line.

A number of factors can impact the determination of the driver status of a non-coding mutation. For example, there are a plethora of ways in which a non-coding mutation may impact genome function. For example, a mutation may alter a transcription factor binding site, affect the partitioning of the genome into topologically-associating domains, or cause epigenetic changes by altering the binding of pioneer factors, nucleosome positioning, chromatin organisation or CpG methylation (1). In this study, we have proposed a list of single nucleotide variants and genomic windows containing recurrent indels, which may be functional mutations in the non-coding genome. We did so by using measures of recurrence, FunSeq2 score (29), and annotations of transcription factor binding. It is possible that others of the recurrent mutations that we identified are actually cancer drivers that impact the genome in a way that is not captured by these analytical methods. It is also possible that many of the mutations that we have selected as potentially functional are actually passenger mutations, and therefore do not act as drivers in colorectal cancer. In our study, we did not find any strong candidate regulatory driver mutations, and so we did not perform any further experimental validation. Ultimately, in order to identify which variants are true cancer driver events, experimental validation of robust putative cancer drivers will be necessary. Currently, experimental validation of this kind is limited by the difficulties involved in designing a cost-effective and high-throughput approach to assess the functional impact of large numbers of non-coding mutations, especially given the many ways in which a mutation may alter gene regulation.

Notably, we did not find any non-coding regions which harboured an excess of functional variants via OncodriveFML (28). Our cohort may be underpowered to detect low frequency driver mutations, which may not significantly stand out from among the background of passenger mutations. Alternatively, poor sequence coverage at some regulatory elements may mean that certain mutations remain undetected. However, it is also possible that the regulatory regions that we sequenced are actually relatively devoid of driver mutations in colorectal cancer, making such events somewhat rare. Interestingly, colorectal cancers do exhibit relatively low numbers of mutations in many regulatory regions such as promoters (39, 40). Mutation loads in colorectal cancer closely follow levels of DNA methylation, and regulatory elements such as these are generally lowly methylated (40).

Since regulatory elements in colorectal cancer accumulate somewhat fewer mutations, it is possible that such regions are subsequently less likely to develop cancer drivers. It may be the case that non-coding driver mutations affecting gene regulation in colorectal cancer are rare in cohorts of this size.

## Conclusions

Taken together, our study has demonstrated TCS to be a sequencing-efficient alternative to traditional WGS analyses when seeking to identify variants at specific loci among larger cohorts. We found that the increased sequencing depth afforded by TCS allows for improved detection of single nucleotide and indel variants, and we demonstrated the utility of TCS for mutational signature analyses. By assessing variant recurrence and function, we proposed some regulatory mutations that may be functional, potentially warranting investigation into whether they play a role in oncogenesis. However, we did not find any strong candidate regulatory driver mutations in the regions that we sequenced, suggesting that with our current sample size, such mutations may be rare.

## Materials and Methods

### Target capture sequencing assay design and analysis of sequencing data

A unique TCS assay was designed to provide sequencing data covering regulatory regions and some coding exons, encompassing almost 36 million nucleotides of the genome (regions listed in Table S1a). Promoter elements were selected to primarily include the region ±450 bp of FANTOM5 p1 promoters of canonical genes (9). DHS sites were selected using HCT-116 DHS sequencing (DNase-seq) hotspot data (Gene Expression Omnibus [GEO] accession: GSM736493). lncRNA, miRNA and DHS sites were prioritised for inclusion into the TCS assay if they were previously recorded to be mutated in other colorectal cancers samples available from TCGA, with further priority given to lncRNAs that were expressed in colon tissue (41). Coding genes included in the TCS assay (Table S1b) are from known colorectal cancer driver genes based in part on gene lists from the COSMIC Cancer Census (20, 21).

95 colorectal cancer and matched normal samples were selected from a pre-existing biobank, and were unbiased for gender, cancer stage or tumour location (**Table 1**, **Table S2**). Fresh tumour tissue had been obtained from surgical resection specimens at St. Vincent’s Hospital, Sydney (ethics numbers H00/022 and 00113). Samples were sequenced using our TCS assay by the Next Generation Sequencing Facility at Western Sydney University, and WGS was additionally performed on a single sample (CRC_1). The TCS was performed using the the Roche NimbleGen SeqCap EZ Exome Library SR platform, version 4.2. The WGS library was prepared with the TruSeq DNA PCR-Free Sample Prep Kit with a 350bp insert size. Both TCS and WGS libraries were sequenced using a 2x101 paired-end read length on the HiSeq 2500. Raw sequencing data has been deposited in European Genomephenome Archive (EGA) under accession number [data deposition in progress].

Raw 101 bp paired-end sequencing reads as fastq files were trimmed using Trim Galore! (https://github.com/FelixKrueger/TrimGalore) to remove 10 bp at the 3' end of reads for the TCS data, and with default parameters for the WGS data. Reads were aligned against assembled chromosomes of hg19 using Burrows-Wheeler Alignment (BWA) mem (42) with default parameters. Files were sorted and indexed with samtools (43) and read groups were added using Picard (https://github.com/broadinstitute/picard). When analysing the WGS data, an additional duplicate removal step was included via the samtools (43) ‘rmdup’ tool with default parameters. Coverage statistics were calculated using samtools (43) ‘depth’ tool across sequenced regions.

Somatic single nucleotide variant calls for TCGA colon cancer samples with WGS were processed as previously described (39) (see **Table S3** for sample names). MSI was designated if the sample was listed as being MSI high (MSI-H) via annotations from TCGA.

### Variant detection and analyses

Germline variants were detected using the GATK pipeline (44), and were visualised in figures using the Integrative Genomics Viewer (IGV) (45, 46) with the BAM files described above. For the identification of somatic single nucleotide and indel mutations, BAM files were additionally filtered to exclude reads which mapped to multiple loci by removing reads marked with the “XA:Z:” and “SA:Z:” flags. Somatic single nucleotide variants were detected with Strelka (10), using the bwa configuration file and default parameters, with the exception of the ‘no depth filters’ option which was selected for analysis of TCS data. VAFs were calculated using bam-readcount (https://github.com/genome/bam-readcount), with default parameters. The violin plot incorporating the VAFs of somatic mutations was created in R using ggplot2 (47). Somatic indels were detected using Strelka (10) with parameters as described above, as well as SvABA (17) and Lancet (18) with default parameters. Segments of assembled chromosomes which had high sequence homology with unplaced scaffolds of hg19 were identified using GMAP (48), and somatic single nucleotide and indel mutations that were within such loci were excluded. Somatic and germline variants were annotated with Annovar (49), to detect any protein-coding alterations.

Mutational signatures (19) were identified through Pearson’s correlation of trinucleotide frequencies in a given sample with those from the COSMIC ‘Signatures of Mutational Processes in Human Cancer’ database (20, 21). Mutational signatures from TCS were normalised against those from the COSMIC database using genome trinucleotide frequencies (“tri.counts.genome”) obtained from the deconstructSigs R package (50). All Pearson’s correlations reported had *P* < 0.0001, indicating a correlation coefficient that is significantly different from zero.

MSI status was determined by analysing mononucleotide repeats, as these sites are error-prone and are typically repaired by the mismatch repair process that becomes deficient in MSI tumours. The mononucleotide markers used were Bat25, Bat26, Bat40 and Cat25, as described previously (11). *POLE* exonuclease domain mutant cancers were identified through manual examination of sequencing data using IGV (45, 46) across the exonuclease domain of *POLE* (amino acids 268-471). This was done for all samples with a somatic exonuclease domain mutation detected by Strelka (10) and/or *r* ≥ 0.75 by Pearson’s correlation with signature 10. (All samples with *r* ≥ 0.75 by Pearson’s correlation with signature 10 did harbour a *POLE* exonuclease domain mutation, and all mutations detected by Strelka (10) were confirmed as somatic via IGV).

### Analysis of regulatory variants for functional or putative driver role

Analyses involving OncodriveFML (version 2.1.0) (28) incorporated both somatic single nucleotide and indel mutations, with ‘targeted’ set as the type of sequencing. The tool was run for coding variants with ‘coding’ set as the type of genomic element (strand provided for coding genes), and was run for all variants with ‘noncoding’ set as the type of genomic element (no strand provided for non-coding regions). All parameters were set to the default, with the exception of the following signatures parameters: method set to ‘bysample’, only_mapped_mutations set to ‘TRUE’ and normalize_by_sites set to “whole_genome”.

FunSeq2 (version 2.1.6) (29) was used to annotate somatic single nucleotide mutations (with no evaluation of recurrence), with the minor allele frequency threshold set to ‘1’ and the maximum length cut-off for indel analyses set to ‘inf’. For variants with different alternate nucleotides between TCS and TCGA cohorts, the alternate nucleotide from the TCS cohort was selected for analysis via FunSeq2. UCSC Genome Browser (51) screenshots show gene predictions via the “UCSC Genes” track. Sequencing data tracks shown in figures have GEO accession numbers as follows: RNA-sequencing (RNA-seq) in HCT-116 cells (GSM958749); H3K4me3 ChIP-seq in HCT-116 cells (GSM945304); DNase-seq in HCT-116 (GSM736600, GSM736493); *SP1* ChIP-seq in HCT116 cells (GSM1010902); and ChIP-seq in HeLa-S3 cells for *E2F4* (GSM935365), *E2F6* (GSM935476) and *MAZ*(GSM935272), for which ChIP-seq data in HCT-116 cells were not available.

### Experimental validation of variants detected

Some somatic mutations were randomly selected for experimental validation via Sanger sequencing of polymerase chain reaction (PCR) product amplified from cancer and matched normal patient DNA. Sanger sequencing was performed by the Ramaciotti Centre for Genomics at the University of New South Wales (UNSW Sydney). Validation was possible for single nucleotide somatic mutations present at > 20% VAF. Mutations at lower VAFs were likely unable to be validated due to the technical limitations of this sequencing method from bulk PCR product. Indels in the putative promoter of *MTERFD3* were also validated as described here.

## Authors’ contributions

Project planning and design: J.E.P., N.H., R.L.W., L.B.H. and J.W.H.W. Experimental analysis: R.C.P., D. Packham and C.J. Data analysis: R.C.P., D. Perera, A.S. and J.W.H.W. Manuscript writing and figures: R.C.P. and J.W.H.W. All authors reviewed and edited the final manuscript.

## Acknowledgements

The authors thank TCGA and other groups who have made their data available for public analysis, and additionally thank Intersect Pty Ltd for providing high-performance computing resources and data storage used in this study.

## Data access

Please contact authors.

## List of Abbreviations

bp: Base pairs
BWA: Burrows Wheeler Aligner
ChIP-seq: Chromatin immunoprecipitation sequencing
COSMIC: Catalogue of Somatic Mutations in Cancer
DHS: DNase I hypersensitivity
DNase-seq: DNase I hypersensitivity sequencing
ENCODE: Encyclopedia of DNA Elements
GEO: Gene Expression Omnibus
IGV: Integrative Genomics Viewer
Indel: Insertion and deletion
lncRNA: Long non-coding RNA
mb: megabase
miRNA: MicroRNA
MSI: Microsatellite instability
MSS: Microsatellite stable
mtDNA: mitochondrial DNA
mTERF: Mitochondrial transcription termination factor
PCR: Polymerase chain reaction
*POLE*: *Polymerase epsilon*
RNA-seq: Ribonucleic acid sequencing
S.D.: Standard deviation
TCGA: The Cancer Genome Atlas
TCS: Target capture sequencing
VAF: Variant allele frequency
WGS: Whole-genome sequencing
WXS: Whole exome sequencing

---

**Figure S1 – Variant allele frequency (VAF) and mutation validation by Sanger sequencing. (a)** Violin plot depicting the VAFs of all single nucleotide somatic variants identified from TCS data (pre-filter) and only variants with VAF >8.5% (VAF-filter). The plot was produced using the ggplot2 R package (47), where the shape indicates the probability density of the data, with mean (dot) and standard deviation (line) indicated. **(b-c)** Sequencing traces from Sanger sequencing of genomic DNA from the samples named, showing validation of **(b)** a somatic deletion and **(c)** four somatic single nucleotide variants. Sequencing traces are visualised using Geneious version 10.2.2 (http://www.geneious.com; (52)).

**Figure S2 – Coverage statistics for whole-genome sequencing (WGS) and target capture sequencing (TCS). (a)** Average per sample TCS read coverage at sequenced bases in cancer (top) and matched normal (bottom) samples. Red bars indicate individual samples sequenced by TCS (*n* = 95). Average coverage across TCS samples is shown by a black dotted line, and average coverage in the WGS sample is shown by a blue dotted line. **(b-f)** Percentage of bases with given read coverage in cancer (top) and matched normal (bottom) samples in **(b)** promoters, **(c)** DNase I hypersensitive (DHS) sites, **(d)** long non-coding RNAs (lncRNAs), **(e)** coding exons and **(f)** microRNAs (miRNAs). Data is plotted in bins spanning 50 reads, where the number on the x-axis indicates the lower edge of the bin (inclusive). The box plot shows the actual value for WGS data (blue; *n* = 1, CRC_1), and the mean and standard deviation across samples in the TCS cohort (red; *n* = 95 samples).

**Figure S3 – Comparison of somatic variants detected from whole-genome sequencing (WGS) and target capture sequencing (TCS) data. (a)** Normalised mutational signatures derived from CRC_1 (top), compared against signature 10 from the COSMIC database (20, 21) (bottom). Signatures are shown for mutations from TCS (left) and WGS (right) data. **(b-c)** Read coverage in cancer and matched normal sequencing data for bases containing somatic variants detected in colorectal cancer sample CRC_1. Graphs show (b) data from TCS for WGS-unique and shared mutations, and **(c)** data from WGS for TCS-unique and shared mutations. Box plots indicate mean and standard deviation of read coverage, where **** denotes *P* < 0.0001.

**Figure S4 – Germline variants and mutational signatures from samples in the target capture sequencing (TCS) cohort.** Snapshot of sequencing reads by TCS from matched normal samples of **(a)** CRC_4 and **(b)** CRC_3. Reads are viewed using the Integrative Genomics Viewer (IGV) (45, 46), with gene transcripts indicated. **(c)** Normalised mutational signatures by TCS (top), against signature 18 from the COSMIC database (20, 21) (bottom) for samples CRC_19 (left), CRC_20 (middle) and CRC_26 (right).

**Figure S5 – Genomic locus harbouring deletions in the MTERFD3 putative promoter, and validation by Sanger sequencing. (a)** Sequencing traces from Sanger sequencing of genomic DNA of the samples named, depicting validation of the three indels within the *MTERFD3* putative promoter. Sequencing traces are visualised using Geneious version 10.2.2 (http://www.geneious.com; (52)). (b) Snapshot from UCSC Genome Browser (51), indicating deletions (indels)within the putative promoter of *MTERFD3*, alongside chromatin immunoprecipitation sequencing (ChIP-seq) data for the transcription factors with motifs disrupted. Boxes contain the reference DNA sequence, with the deleted nucleotides marked by an orange box. Transcription factor binding motifs are shown from Factorbook (31), where a green bar depicts the span of the motif across the DNA sequence.

## References

1. Poulos RC, Wong JWH. cis-Regulatory Driver Mutations in Cancer Genomes. eLS: John Wiley & Sons, Ltd; 2017. p. 1–10.

2. Melton C, Reuter JA, Spacek DV, Snyder M. Recurrent somatic mutations in regulatory regions of human cancer genomes. Nat Genet. 2015;47(7):710–6.

3. Fredriksson NJ, Ny L, Nilsson JA, Larsson E. Systematic analysis of noncoding somatic mutations and gene expression alterations across 14 tumor types. Nat Genet. 2014;46:1258–63.

4. Weinhold N, Jacobsen A, Schultz N, Sander C, Lee W. Genome-wide analysis of noncoding regulatory mutations in cancer. Nat Genet. 2014;46.

5. Rheinbay E, Parasuraman P, Grimsby J, Tiao G, Engreitz JM, Kim J, et al. Recurrent and functional regulatory mutations in breast cancer. Nature. 2017;547(7661):55–60.

6. Lawrence MS, Stojanov P, Mermel CH, Robinson JT, Garraway LA, Golub TR. Discovery and saturation analysis of cancer genes across 21 tumour types. Nature. 2014;505:495–501.

7. Lawrence MS, Stojanov P, Polak P, Kryukov GV, Cibulskis K, Sivachenko A, et al. Mutational heterogeneity in cancer and the search for new cancer-associated genes. Nature. 2013;499(7457):214–8.

8. Guo Y, Long J, He J, Li C-l, Cai Q, Shu X-O, et al. Exome sequencing generates high quality data in non-target regions. BMC genomics. 2012;13(1):194.

9. The FANTOM Consortium and the RIKEN PMI and CLST (DGT). A promoter-level mammalian expression atlas. Nature. 2014;507(7493):462–70.

10. Saunders CT, Wong WS, Swamy S, Becq J, Murray LJ, Cheetham RK. Strelka: accurate somatic small-variant calling from sequenced tumor-normal sample pairs. Bioinformatics. 2012;28(14):1811–7.

11. Hawkins NJ, Ward RL. Sporadic colorectal cancers with microsatellite instability and their possible origin in hyperplastic polyps and serrated adenomas. Journal of the National Cancer Institute. 2001;93(17): 1307–13.

12. Rayner E, van Gool LC, Palles C, Kearsey SE, Bosse T, Tomlinson I, et al. A panoply of errors: polymerase proofreading domain mutations in cancer. Nature reviews Cancer. 2016;16(2):71–81.

13. The Cancer Genome Atlas Research Network. Comprehensive molecular characterization of human colon and rectal cancer. Nature. 2012;487(7407):330–7.

14. Rowan AJ, Lamlum H, Ilyas M, Wheeler J, Straub J, Papadopoulou A, et al. APC mutations in sporadic colorectal tumors: A mutational “hotspot” and interdependence of the “two hits”. Proc Natl Acad Sci USA. 2000;97(7):3352–7.

15. Shen L, Toyota M, Kondo Y, Lin E, Zhang L, Guo Y, et al. Integrated genetic and epigenetic analysis identifies three different subclasses of colon cancer. Proc Natl Acad Sci USA. 2007; 104(47): 18654–9.

16. Rajagopalan H, Bardelli A, Lengauer C, Kinzler KW, Vogelstein B, Velculescu VE. Tumorigenesis: RAF/RAS oncogenes and mismatch-repair status. Nature. 2002;418(6901):934.

17. Wala J, Bandopadhayay P, Greenwald N, O'Rourke R, Sharpe T, Stewart C, et al. Genomewide detection of structural variants and indels by local assembly. bioRxiv. 2017:https://doi.org/10.1101/105080.

18. Narzisi G, Corvelo A, Arora K, Bergmann E, Shah M, Musunuri R, et al. Lancet: genome-wide somatic variant calling using localized colored DeBruijn graphs. bioRxiv. 2017:https://doi.org/10.1101/196311.

19. Alexandrov LB, Nik-Zainal S, Wedge DC; Aparicio SAJR; Behjati S, Biankin AV, et al. Signatures of mutational processes in human cancer. Nature. 2013;500(7463):415–21.

20. Forbes SA; Beare D; Gunasekaran P, Leung K; Bindal N; Boutselakis H; et al. COSMIC: exploring the world's knowledge of somatic mutations in human cancer. Nucleic Acids Res. 2015;43(Database issue):D805–11.

21. Forbes SA, Bindal N, Bamford S, Cole C, Kok CY, Beare D, et al. COSMIC: mining complete cancer genomes in the Catalogue of Somatic Mutations in Cancer. Nucleic Acids Res. 2011;39:D945–50.

22. Plazzer JP, Sijmons RH, Woods MO, Peltomaki P, Thompson B, Den Dunnen JT, et al. The InSiGHT database: utilizing 100 years of insights into Lynch syndrome. Familial cancer. 2013;12(2):175–80.

23. Deng G, Bell I, Crawley S, Gum J, Terdiman JP, Allen BA, et al. BRAF mutation is frequently present in sporadic colorectal cancer with methylated hMLH1, but not in hereditary nonpolyposis colorectal cancer. Clinical cancer research : an official journal of the American Association for Cancer Research. 2004;10(1 Pt 1):191–5.

24. Pilati C, Shinde J, Alexandrov LB, Assié G, André T, Hélias-Rodzewicz Z, et al. Mutational signature analysis identifies MUTYH deficiency in colorectal cancers and adrenocortical carcinomas. The Journal of pathology. 2017;242(1):10–5.

25. Lek M, Karczewski KJ, Minikel EV, Samocha KE, Banks E, Fennell T, et al. Analysis of proteincoding genetic variation in 60,706 humans. Nature. 2016;536(7616):285–91.

26. Ali M, Kim H, Cleary S, Cupples C, Gallinger S, Bristow R. Characterization of mutant MUTYH proteins associated with familial colorectal cancer. Gastroenterology. 2008;135(2):499–507.

27. Chang J, Tan W, Ling Z, Xi R, Shao M, Chen M, et al. Genomic analysis of oesophageal squamous-cell carcinoma identifies alcohol drinking-related mutation signature and genomic alterations. Nat Commun. 2017;8:15290.

28. Mularoni L, Sabarinathan R, Deu-Pons J, Gonzalez-Perez A, López-Bigas N. OncodriveFML: a general framework to identify coding and non-coding regions with cancer driver mutations. Genome biology. 2016;17(1):128.

29. Fu Y, Liu Z, Lou S, Bedford J, Mu XJ, Yip KY, et al. FunSeq2: a framework for prioritizing noncoding regulatory variants in cancer. Genome biology. 2014;15(10):480.

30. Saadeh H, Schulz R. Protection of CpG islands against de novo DNA méthylation during oogenesis is associated with the recognition site of E2f1 and E2f2. Epigenetics Chromatin. 2014;7:26-.

31. Wang J, Zhuang J, Iyer S, Lin XY, Greven MC, Kim BH, et al. Factorbook.org: a Wiki-based database for transcription factor-binding data generated by the ENCODE consortium. Nucleic Acids Res. 2013;41(Database issue):D171–6.

32. The Encode Project Consortium. An Integrated Encyclopedia of DNA Elements in the Human Genome. Nature. 2012;489(7414):57–74.

33. Kloor M, Michel S, Buckowitz B, Ruschoff J, Buttner R, Holinski-Feder E, et al. Beta2-microglobulin mutations in microsatellite unstable colorectal tumors. Int J Cancer. 2007;121(2):454–8.

34. Hyvarinen AK, Pohjoismaki JL, Holt IJ, Jacobs HT. Overexpression of MTERFD1 or MTERFD3 impairs the completion of mitochondrial DNA replication. Molecular biology reports. 2011;38(2):1321–8.

35. Reznik E, Miller ML, Senbabaoglu Y, Riaz N, Sarungbam J, Tickoo SK, et al. Mitochondrial DNA copy number variation across human cancers. eLife. 2016;5:e10769.

36. Mansour MR, Abraham BJ, Anders L, Berezovskaya A, Gutierrez A, Durbin AD, et al. Oncogene regulation. An oncogenic super-enhancer formed through somatic mutation of a noncoding intergenic element. Science. 2014;346(6215):1373–7.

37. Rahman S, Magnussen M, León TE, Farah N, Li Z, Abraham BJ, et al. Activation of the LM02 oncogene through a somatically acquired neomorphic promoter in T-cell acute lymphoblastic leukemia. Blood. 2017;129(24):3221–6.

38. Abraham BJ, Hnisz D, Weintraub AS, Kwiatkowski N, Li CH, Li Z, et al. Small genomic insertions form enhancers that misregulate oncogenes. Nat Commun. 2017;8:14385.

39. Perera D, Poulos RC, Shah A, Beck D, Pimanda JE, Wong JWH. Differential DNA repair underlies mutation hotspots at active promoters in cancer genomes. Nature. 2016;532(7598):259–63.

40. Poulos RC, Olivier J, Wong JWH. The interaction between cytosine méthylation and processes of DNA replication and repair shape the mutational landscape of cancer genomes. Nucleic Acids Res. 2017;45(13):7786–95.

41. Cabili MN, Trapnell C, Goff L, Koziol M, Tazon-Vega B, Regev A, et al. Integrative annotation of human large intergenic noncoding RNAs reveals global properties and specific subclasses. Genes Dev. 2011;25:1915–27.

42. Li H, Durbin R. Fast and accurate short read alignment with Burrows-Wheeler transform. Bioinformatics. 2009;25(14): 1754–60.

43. Li H. The Sequence Alignment/Map format and SAMtools. Bioinformatics. 2009;25:2078–2-9.

44. McKenna A. The Genome Analysis Toolkit: a MapReduce framework for analyzing next-generation DNA sequencing data. Genome research. 2010;20:1297–303.

45. Robinson JT, Thorvaldsdottir H, Winckler W, Guttman M, Lander ES, Getz G, et al. Integrative genomics viewer. Nat Biotech. 2011;29(1):24–6.

46. Thorvaldsdóttir H, Robinson JT, Mesirov JP. Integrative Genomics Viewer (IGV): highperformance genomics data visualization and exploration. Brief Bioinform. 2013;14(2):178–92.

47. Wickham H. ggplot2: Elegant Graphics for Data Analysis. New York: Springer International Publishing; 2016.

48. Wu TD, Watanabe CK. GMAP: a genomic mapping and alignment program for mRNA and EST sequences. Bioinformatics. 2005;21(9):1859–75.

49. Wang K, Li M, Hakonarson H. ANNOVAR: functional annotation of genetic variants from highthroughput sequencing data. Nucleic Acids Res. 2010;38(16):e164–e.

50. Rosenthal R, McGranahan N, Herrero J, Taylor BS, Swanton C. deconstructSigs: delineating mutational processes in single tumors distinguishes DNA repair deficiencies and patterns of carcinoma evolution. Genome biology. 2016;17:31.

51. Kent WJ, Sugnet CW, Furey TS, Roskin KM, Pringle TH, Zahler AM, et al. The human genome browser at UCSC. Genome research. 2002;12(6):996–1006.

52. Kearse M, Moir R, Wilson A, Stones-Havas S, Cheung M, Sturrock S, et al. Geneious Basic: an integrated and extendable desktop software platform for the organization and analysis of sequence data. Bioinformatics. 2012;28(12):1647–9.

